# Targeted mRNA delivery with bispecific antibodies that tether LNPs to cell-surface markers

**DOI:** 10.1101/2024.10.17.618962

**Authors:** Bettina Dietmair, James Humphries, Timothy R. Mercer, Kristofer J. Thurecht, Christopher B. Howard, Seth W. Cheetham

**Affiliations:** Australian Institute for Bioengineering and Nanotechnology, The University of Queensland, Brisbane, QLD, Australia; BASE facility, The University of Queensland, Brisbane, QLD, Australia; Centre for Advanced Imaging, ARC Research Hub for Advanced Manufacture of Targeted Radiopharmaceuticals, The University of Queensland, St Lucia, QLD, 4072, Australia

## Abstract

Efficient delivery of mRNA-LNPs to specific cell-types remains a major challenge in the widespread application of mRNA therapeutics. Conventional targeting approaches involve modifying the lipid composition or functionalising the surface of lipid nanoparticles (LNPs), which complicates manufacturing, alters nanoparticle size, charge and stealth, impacting their delivery and immunogenicity. Here we present a generalisable method for targeted mRNA-LNP delivery that uses bispecific antibodies (BsAbs) to form a bridge between LNPs and cell-surface markers. Instead of attaching the targeting agent to the nanocarrier, BsAbs are administered first, bind to surface proteins on target cells, and later retain unmodified LNPs in affected tissues. We demonstrate efficient and cell-type-specific delivery of mRNA-LNPs to epidermal growth factor receptor (EGFR), and folate hydrolase 1 (PSMA) positive cells *in vitro* and *in vivo*. The flexibility of this technology, achieved by substitution of the cell-targeting region of the BsAbs, enables rapid development of next-generation targeted mRNA drugs.

## Main

mRNA therapies are emerging rapidly as a new class of drugs with the potential to treat a wide range of human diseases. Beyond vaccines, mRNA applications include the treatment of cancers^1^, autoimmunity^2^, and hereditary diseases^3^. The efficacy of mRNA drugs depends on the ability to deliver mRNA to specific cell-types at sufficient dosage. Formulation into LNPs protects the mRNA during storage and circulation, and enables delivery of the negatively charged nucleic acid across the cell membrane^4^. Polyethylene glycol (PEG) coating of the LNP surface confers “stealth” properties by forming a hydration layer that stabilises LNPs and decreases protein interactions, thus reducing immune recognition^5^. PEGylated LNPs are clinically validated as safe and effective mRNA delivery systems^6,7,8^. However, following intravenous administration, LNP accumulation and mRNA expression occurs mainly in the liver^9,10.^ Hepatic uptake is promoted by the lipid constitution of LNPs^11^, the size of the nanoparticles allowing passage through liver fenestrae^12^, and protein adsorption to the LNP surface enhancing receptor-mediated hepatocyte uptake^13^. The liver tropism of LNPs limits the potential of mRNA for treating extrahepatic disease.

Local administration to extrahepatic tissues is unsuitable for broad application as it is invasive, and challenging for dispersed target cells, such as immune cells, stem cells and tumour metastases^14^. Systemic administration is therefore preferred but requires efficient and cell-specific targeting of mRNA-LNPs. Passive targeting harnesses enhanced vascular permeability and reduced lymphatic drainage around pathological tissues for increased nanoparticle access and retention^15^. Extended LNP circulation achieved by PEGylation enables nanoparticles to reach peripheral tissues but passive delivery is inconsistent and inefficient^5,16.^ To actively target LNPs to specific cell types, intensive efforts have focused on modifying the lipid composition of LNPs^17^ or functionalising the surface with targeting moieties, including ligands and antibody fragments^18^. Lipid conjugates complicate manufacturing of LNPs, hindering scale-up production and clinical translation^19^. Incorporation of modifications during LNP production can result in encapsulation of the targeting ligand, reducing the availability of target-binding sites on the nanoparticle surface. Post-synthesis LNP functionalisation can increase accessibility of the binding moiety, though attachment can still occur in the wrong orientation^20^. Although these LNP modifications can improve target accumulation and receptor-mediated internalisation, they shift LNP charge, increase the nanoparticle size, and change the protein adsorption profile, leading to altered biodistribution, uptake and immunogenicity of the nanocarrier^21^.

Here we describe a customisable method that uses bispecific antibodies for targeted mRNA-LNP delivery. The BsAbs comprise two linked single-chain variable fragments (scFvs) that can bind to PEG on the exterior of LNPs and to a protein enriched on the target cell surface. Cell-specific mRNA-LNP delivery can be achieved by non-covalent attachment of BsAbs to the surface of the LNPs before administration (pre-mixing; **Fig. 1a**) or by pre-targeting (**Fig. 1b**), whereby cells are initially exposed to BsAbs, followed by administration of unmodified LNPs^22^. Targeting with BsAbs enabled mRNA drug delivery beyond the liver, efficiently increasing mRNA uptake and expression in specific cell-types and decreasing uptake in off-target tissues. Separate administration of BsAbs and mRNA-LNPs significantly outperformed targeted delivery using pre-mixed mRNA-LNPs and BsAbs, demonstrating specificity for EGFR and PSMA *in vitro* and *in vivo*. This approach has the potential to significantly broaden the reach of mRNA medicines.

**Fig 1:**
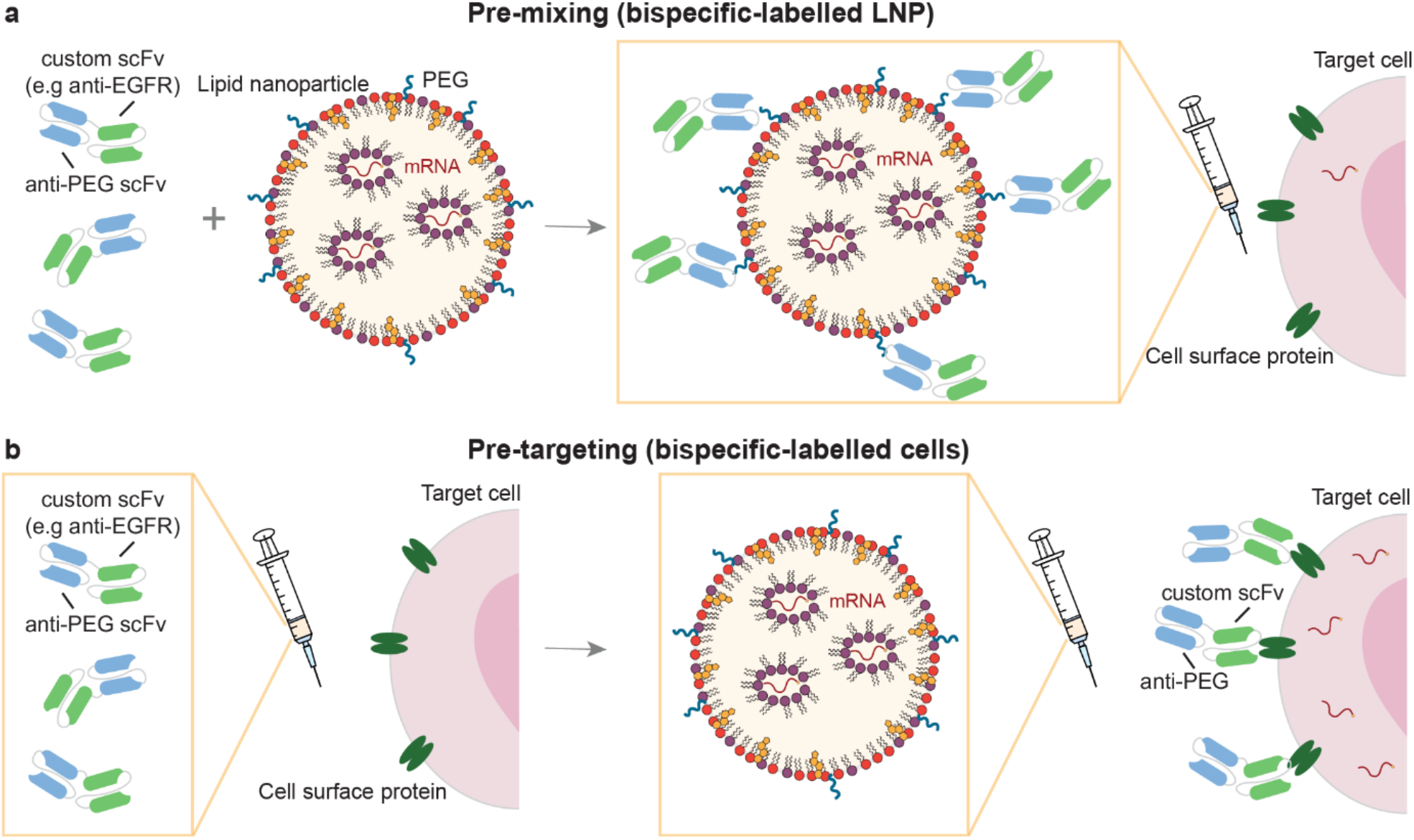
Bispecific antibodies enable cell-specific mRNA-LNP delivery. **a**, During pre-mixing, bispecific antibodies (BsAbs) bind to polyethylene glycol (PEG) on the surface of mRNA-loaded lipid nanoparticles (LNPs). The second binding region of the BsAb can bind to surface protein on the target cell. **b**, For pre-targeting, cells are exposed to BsAbs that can specifically bind to surface proteins. mRNA-carrying LNPs can then bind to the PEG-specific binding region of the BsAb.

### Pre-targeting with BsAbs improves cell-specific mRNA-LNP delivery

To determine if cell-surface antigen-specific BsAbs enable targeted mRNA-LNP delivery, we synthesised and encapsulated enhanced green fluorescent protein (eGFP) mRNA in LNPs (**Supplementary fig. 1a**). As active targeting is commonly achieved by attaching the targeting agent to the LNPs (pre-mixing), not to the target cells (pre-targeting), we first characterised the effect of surface functionalisation on LNP properties. Pre-mixing of the mRNA-LNPs with BsAbs increased the median size of the mRNA-LNP from 82 nm to 181 nm and 412 nm for anti-PSMA and anti-EGFR BsAbs, respectively (**Fig. 2a**). BsAb-coating altered LNP morphology, reduced particle uniformity and induced aggregation (**Fig. 2a,b**), which may contribute to LNP behaviour *in vivo*^23,24^. *For intravenous administration and tissue penetration nanoparticle sizes below 150 nm are preferred*^*25*^.

**Fig 2:**
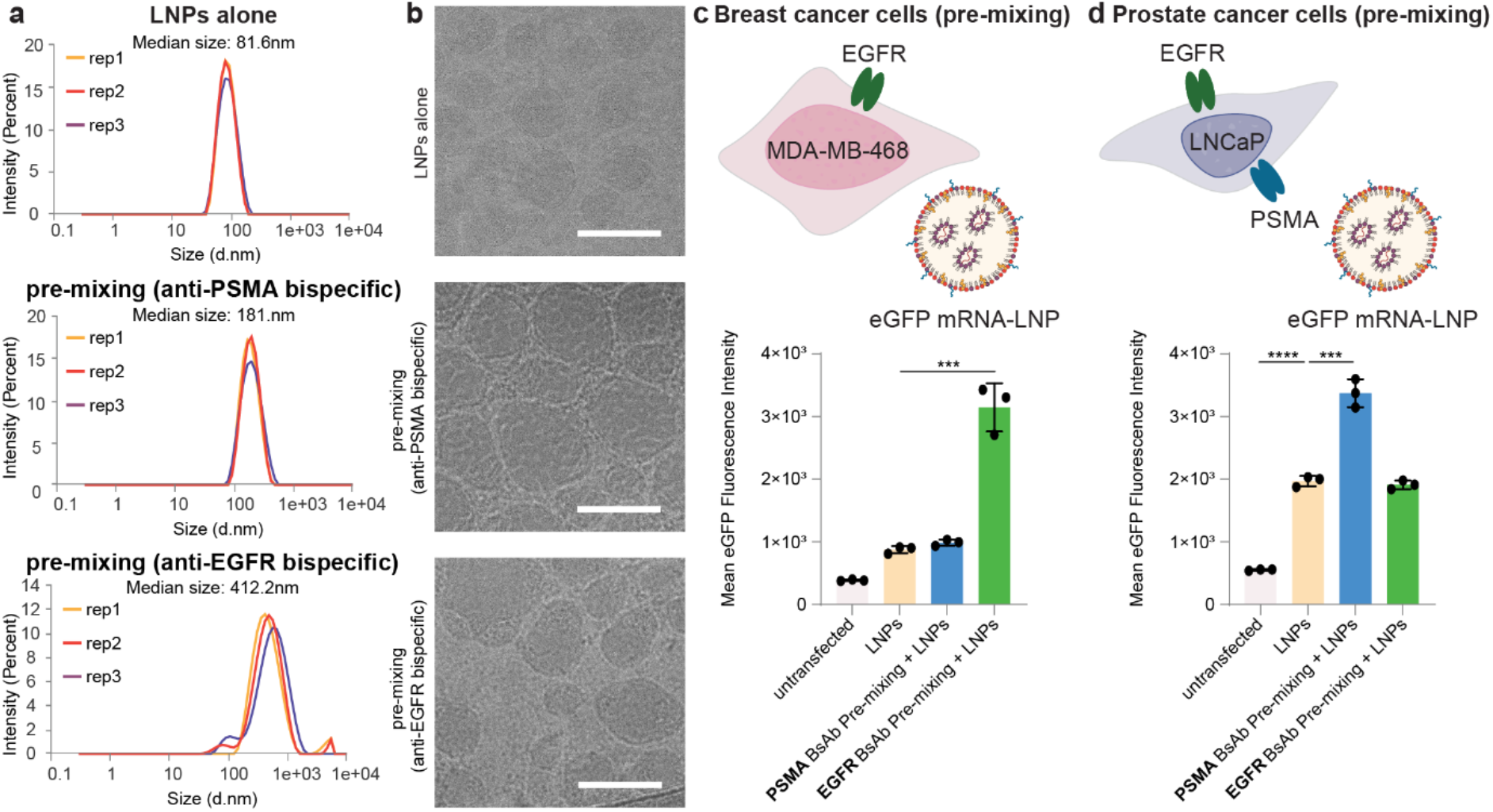
Pre-mixing of cells with bispecific antibodies alters physicochemical properties and delivery of mRNA-LNPs. **a**, Triplicate dynamic light scattering measurements of size distribution of eGFP-mRNA LNPs without BsAbs, after pre-mixing with PSMA-PEG BsAbs and after pre-mixing with EGFR-PEG BsAbs. **b**, Cryo-transmission electron microscopy images of eGFP-mRNA LNPs without BsAbs, after pre-mixing with PSMA-PEG BsAbs and after pre-mixing with EGFR-PEG BsAbs. Scale bar: 100 nm. **c**, Mean eGFP fluorescence intensity of cells transfected with untargeted eGFP-mRNA LNPs, with LNPs pre-mixed with PSMA-PEG BsAbs or LNPs pre-mixed with EGFR-PEG BsAbs, respectively, for MDA-MB-468 breast cancer cells (PSMA-ve, EGFR+ve) and **d**) LNCaP prostate cancer cells (PSMA+ve, EGFR+ve). Mean eGFP fluorescence intensity was measured using flow cytometry. Statistical analysis was performed using two-tailed t-tests assuming equal variance. Bars represent the mean value, error bars indicate standard deviation (n = 3). *p < 0.05, **p < 0.01, ***p < 0.001 and ∗∗∗∗p < 0.0001.

To evaluate cell-type-specific delivery of mRNA-LNPs pre-mixed with BsAbs, we used MDA-MB-468 breast cancer cells which express the epidermal growth factor receptor (EGFR), but not folate hydrolase 1 (PSMA), and LNCaP prostate cancer cells which express both EGFR and PSMA (**Supplementary fig. 1b**). Pre-mixing of LNPs with anti-PEG:anti-EGFR BsAbs enhanced mRNA-LNP delivery to MDA-MB-468 (**Fig. 2c**). LNP surface functionalisation with anti-PEG:anti-PSMA BsAbs did not improve mRNA-LNP delivery to the PSMA-negative cell line. To confirm cell-type-specificity of BsAb-mediated targeting, we analysed mRNA-LNP delivery to EGFR+ve, PSMA+ve LNCaP prostate cancer cells (**Fig. 2d**). eGFP mRNA-LNP delivery significantly improved when mRNA-LNPs were pre-mixed with anti-PEG:anti-PSMA BsAbs. Pre-mixing with anti-PEG:anti-EGFR BsAbs had no effect on eGFP expression compared to untargeted LNPs.

We then compared the established pre-mixing protocol to pre-targeting of cells with BsAbs. Pre-targeting of MDA-MB-468 breast cancer cells with anti-PEG:anti-EGFR BsAbs significantly increased eGFP expression compared to pre-mixing (twelve-fold vs. three-fold improvement over mRNA-LNPs alone; **Fig. 3a,b, Supplementary fig. 1c**). Similarly, pre-targeting of LNCaP prostate cancer cells reached almost three-fold improvement with PSMA BsAbs and four-fold improvement with EGFR BsAbs, compared to pre-mixing (**Fig. 3c,d, Supplementary fig. 1d**). Compared to untargeted LNP transfection, pre-targeting with EGFR-PEG and PSMA-PEG achieved four-fold and five-fold eGFP expression, respectively. Notably, pre-targeting with EGFR-PEG BsAbs enhanced mRNA-LNP delivery to EGFR+ve LNCaP cells while pre-mixing did not improve uptake.

**Fig 3:**
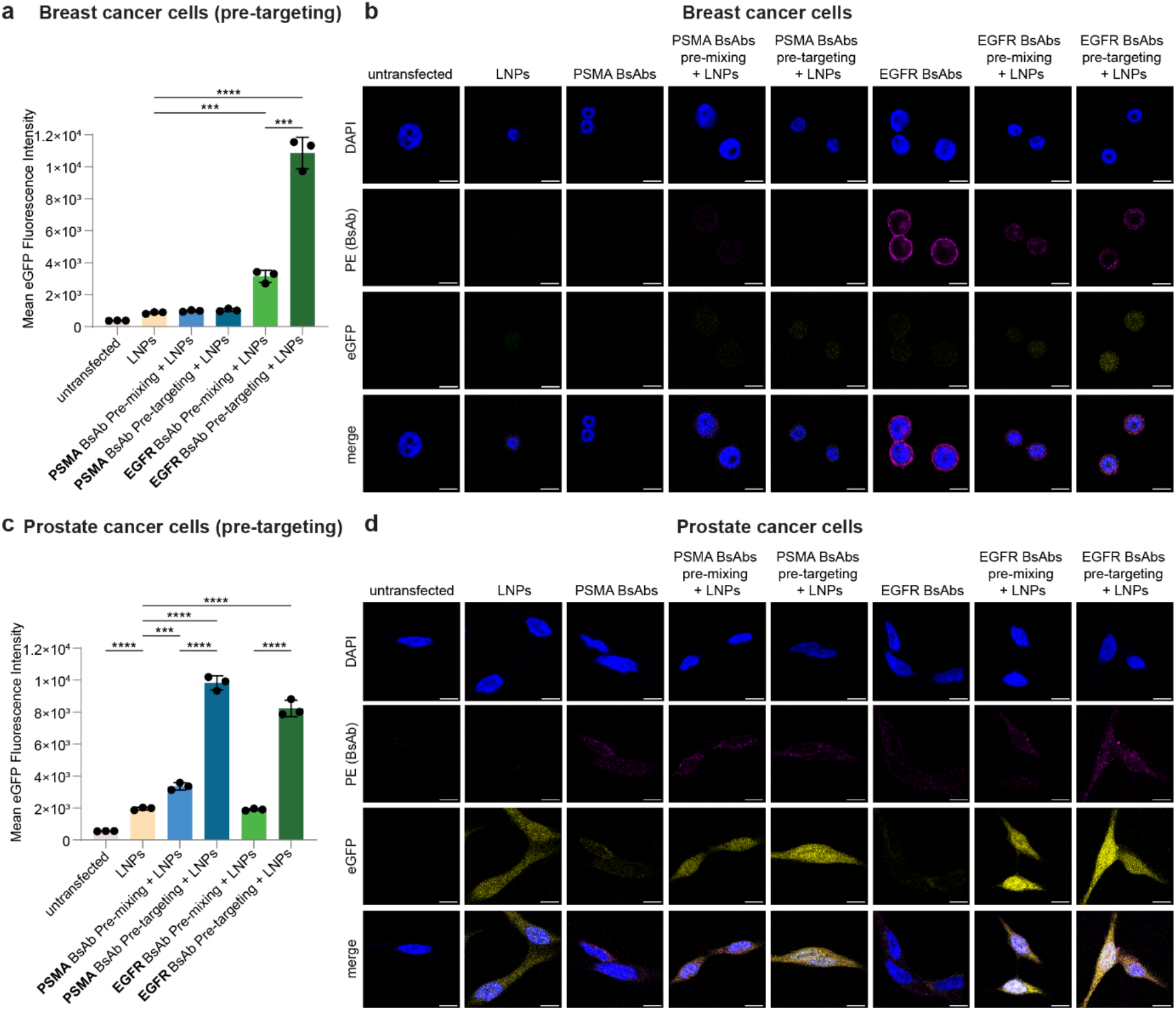
Pre-targeting of cells with bispecific antibodies improves cell-specific delivery of mRNA-LNPs. **a**, Mean eGFP fluorescence intensity of MDA-MB-468 breast cancer cells (PSMA-ve, EGFR+ve) transfected with eGFP-mRNA LNPs. PSMA-PEG BsAbs or EGFR-PEG BsAbs were pre-mixed with LNPs or pre-targeted to MDA-MB-468 cells, respectively. **b**, Confocal microscopy images of MDA-MB-468 after addition of eGFP mRNA-LNPs, PSMA-PEG or EGFR-PEG BsAbs, BsAbs pre-mixed with LNPs or pre-targeting with BsAbs followed by addition of LNPs. **c**, Mean eGFP fluorescence intensity of LNCaP prostate cancer cells (PSMA+ve, EGFR+ve) transfected with eGFP-mRNA LNPs. PSMA-PEG BsAbs or EGFR-PEG BsAbs were pre-mixed with LNPs or pre-targeted to LNCaP cells, respectively. **d**, Confocal microscopy images of LNCaP after addition of eGFP mRNA-LNPs, PSMA-PEG or EGFR-PEG BsAbs, BsAbs pre-mixed with LNPs or pre-targeting with BsAbs followed by addition of LNPs. Confocal microscopy images were taken at 63 × magnification and show eGFP expression (yellow), BsAb localisation (protein L-phycoerythrin conjugate labelled; magenta) and 4′,6-diamidino-2-phenylindole DNA stain (DAPI; blue) four hours after addition of LNPs. Scale bar: 10 μm. Mean eGFP fluorescence intensity was measured using flow cytometry. Statistical analysis was performed using two-tailed t-tests assuming equal variance. Bars represent the mean value, error bars indicate standard deviation (n = 3). *p < 0.05, **p < 0.01, ***p < 0.001 and ∗∗∗∗p < 0.0001.

In summary, pre-mixing and pre-targeting with anti-PEG BsAbs facilitates cell-type-specific mRNA-LNP delivery *in vitro*. The pre-targeting approach is applicable to different cell lines and target antigens and is more efficient than pre-mixing. Active targeting employs interactions with a cell surface receptor to promote specificity and internalisation *via* endocytosis^26^. As BsAbs are below the size range in which receptor-mediated endocytosis is triggered^27,28,^ they accumulate on the plasma membrane and are internalised once mRNA-LNPs bind (**Fig. 3b,d**). In addition to maintaining the physicochemical properties of mRNA-LNPs by pre-targeting, avidity might improve delivery with more BsAb binding sites available on cell surfaces during pre-targeting than PEG on LNPs during pre-mixing. As pre-targeting of MDA-MB-468 with EGFR-PEG BsAbs achieved the greatest improvement over untargeted and pre-mixed LNP delivery *in vitro*, we tested this condition *in vivo*.

### Pre-targeting with BsAbs improves targeted mRNA-LNP delivery *in vivo*

To evaluate the efficacy of BsAb-targeted mRNA-LNP delivery *in vivo*, we synthesised and encapsulated firefly luciferase mRNA in LNPs (**Supplementary fig. 2a,b,c**) and pre-mixed or pre-targeted with EGFR-PEG BsAbs for intravenous administration to BALB/c nude mice with subcutaneous MDA-MB-468 xenografts (**Fig. 4a**). Pre-mixing and pre-targeting with EGFR-PEG BsAbs increased the mRNA-LNP delivery to the tumour tissue over eight-fold and seven-fold while reducing the radiance in the liver by a third and half compared to untargeted LNPs, respectively, eight hours after luciferase mRNA-LNP administration (**Fig. 4b**). After 48 hours, luciferase expression in the liver was reduced across all delivery groups (**Fig. 4c**). In contrast, luminescence in the tumour remained consistent with the eight-hour timepoint in pre-targeted animals, only decreasing by around ten percent. Radiance in the pre-mixing group tumours was reduced by 60 percent compared to the measurement after eight hours.

**Fig 4:**
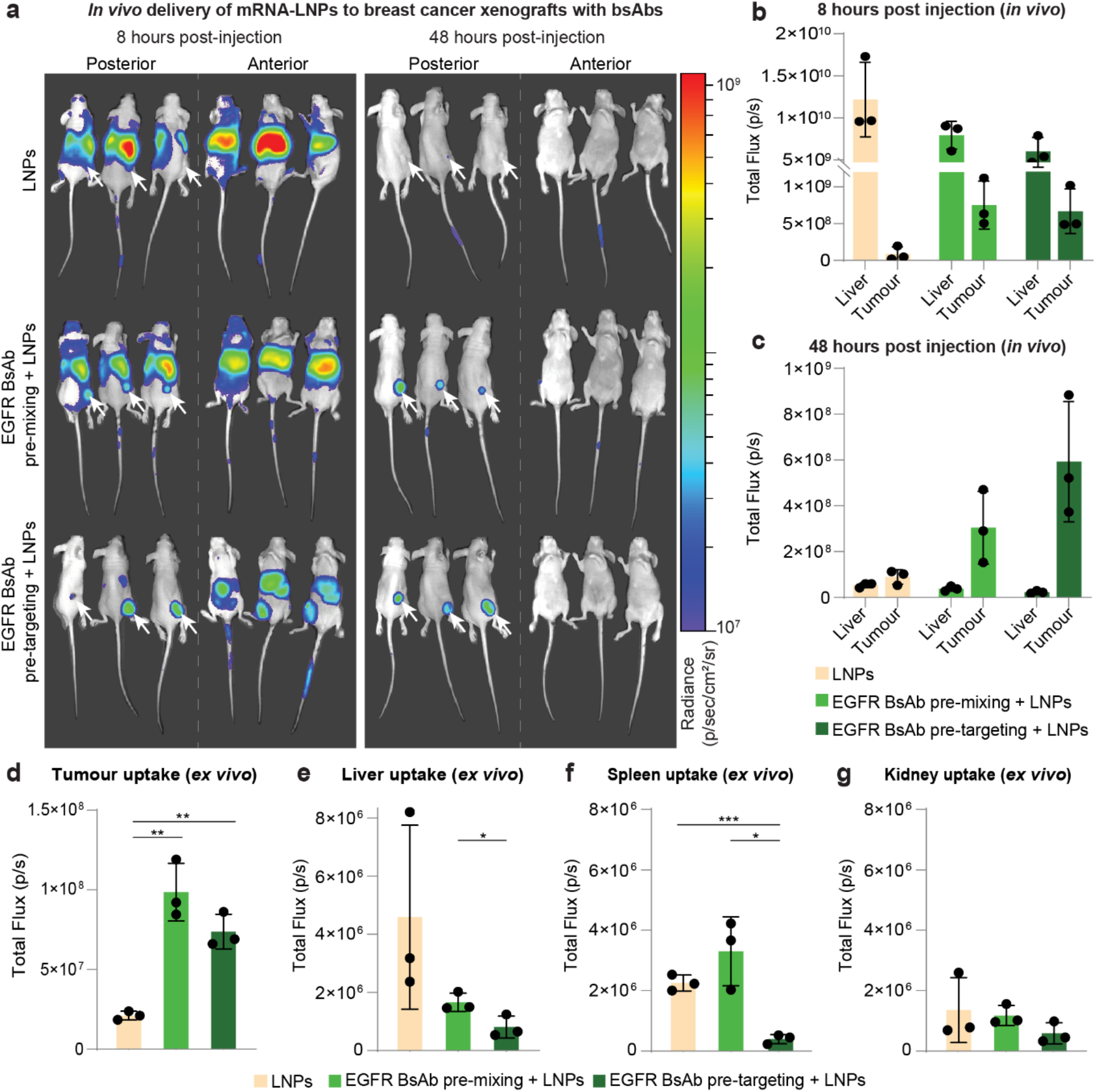
Pre-targeting with bispecific antibodies improves targeted delivery of mRNA-LNPs in vivo. **a**, In vivo bioluminescence images of MDA-MB-468 tumour-bearing mice injected with untargeted luciferase mRNA-LNPs, LNPs pre-mixed with EGFR-PEG BsAbs or mice pre-injected with EGFR-PEG BsAbs followed by administration of mRNA-LNPs. White arrows indicate posterior tumour localisation. **b**, In vivo bioluminescence in the liver compared to tumour for different targeting approaches eight hours and **c**, 48 hours after luciferase mRNA-LNP administration **d**, Ex vivo bioluminescence imaging of tumour, **e**, liver, **f**, spleen and **g**, kidney tissue at 48 hours post-injection. Background was subtracted based on a saline-injected mouse. Statistical analysis was performed using two-tailed t-tests assuming equal variance. Bars represent the mean value, error bars indicate standard deviation (n = 3), *p < 0.05, **p < 0.01 and ***p < 0.001.

*Ex vivo* analysis of tumour signal confirmed *in vivo* biodistribution results, showing comparable luminescence between pre-mixing and pre-targeting cohorts with more than 4.6-fold and 3.5-fold elevated radiance compared to untargeted mRNA-LNP administration, respectively (**Fig. 4d**). Consistently, total flux in the liver was highest in the untargeted LNP group and significantly lower in mice treated with pre-targeting, compared to pre-mixing of BsAbs and LNPs (**Fig. 4e**). Likewise, spleen uptake was lowest in pre-targeted mice compared to over five-fold and eight-fold increased spleen signal for administration of untargeted and pre-mixed mRNA-LNPs, respectively (**Fig. 4f**). High splenic uptake of LNPs pre-mixed with BsAbs could result from changes in physicochemical properties, including a large protein corona around the LNPs^24,29^ and a charge shift from positively charged, unmodified LNPs to negatively charged, BsAb pre-mixed LNPs^16^ (**Supplementary fig. 2d**). Radiance was low in the heart (**Supplementary fig. 2e**), blood (**Supplementary fig. 2f**) and the second major clearance organ, kidney (**Fig. 4g**).

Confirming the *in vitro* results, targeting mRNA-LNPs with EGFR-PEG BsAbs *in vivo* resulted in mRNA delivery to EGFR+ve tumours and reduced delivery to the liver, demonstrating specificity and efficacy compared to delivery of untargeted LNPs. Pre-targeting led to sustained mRNA expression in the tumour and lowest signal in the liver and spleen compared to pre-mixing of LNPs with BsAbs or administration of untargeted LNPs.

## Conclusions

We demonstrated efficient mRNA-LNP targeting *in vitro* and *in vivo* using BsAbs specific to LNPs and cell-surface markers. Temporal separation of BsAb and nanoparticle administration *via* pre-targeting achieved superior mRNA uptake and expression. Pre-targeting could improve biodistribution of mRNA-LNPs relative to pre-mixed systems by reducing the effect of protein fouling on the particle surface^22^. Lower mRNA dosage and reduced uptake in off-target organs, such as the liver and spleen, could reduce the toxicity of mRNA therapies. While mRNA-LNPs were cleared from the metabolically active liver^5^, efficient mRNA expression persisted exclusively in the tumour. BsAbs enable mRNA delivery to previously inaccessible tissues which broadens the scope of conditions addressable with mRNA therapeutics. To the best of our knowledge, this is the first time pre-targeting has been characterised for mRNA-LNPs. The structure of the BsAbs facilitates substitution of the antigen-specific binding domain, enabling customised targeting of unmodified, PEG-containing mRNA carriers to markers present on individual patient cell types. Separate production of the mRNA-LNP and targeting agent (BsAb) greatly streamlines manufacture compared to functionalised-mRNA-LNPs^30^. The well-established clinical use and safety of mRNA-LNPs^31^ and BsAbs^32^ provides strong precedence for a streamlined clinical translation of this technology. Efficient and adaptable targeting of mRNA-LNPs enables rapid development of next-generation mRNA drugs, including protein replacement therapies and gene editing applications for incurable disease.

## Methods

### Template production

eGFP and firefly luciferase mRNA templates were designed with the CleanCap® AG promoter^33^, human alpha-globin 5′ UTR and the mouse alpha-globin 3′ UTR sequences and synthesised as gBlocks™ HiFi Gene Fragments (IDT™). 0.5 ng of Gene Fragments, forward primer (10 μM), reverse primer (10 μM), and NEB® Q5® HotStart 2 × master mix were combined at room temperature in a 100 μL PCR reaction and placed on ice. The template was amplified using the following thermocycler program.

| Step | Cycle | Temperature | Time |
| --- | --- | --- | --- |
| Denaturation | 20 × | 98 °C | 10 seconds |
| Annealing |  | 64 °C | 30 seconds |
| Extension |  | 72 °C | 30 seconds per kb |
| Final extension | 1 × | 72 °C | 2 minutes |
| Hold | 1 × | 4 °C | Indefinite |

Eight PCR reactions were purified using the Qiagen® QIAquick® PCR cleanup kit, according to the manufacturer’s instructions. The PCR product was eluted in 30 μL of ultrapure water, analysed by gel electrophoresis and quantified using UV spectrophotometry.

### mRNA production

Amplified Gene Fragments were used as template for mRNA *in vitro* transcription (IVT) at a concentration of 50 μg/mL. IVT was performed using 16 μg/mL T7 RNA polymerase (New England Biolabs; NEB® M0251), ribonucleotides (6 mM ATP, 5 mM CTP, 5 mM GTP; NEB®), 5 mM N1-methylpseudouridine-5’-triphosphate (TriLink® BioTechnologies, TRN1081), 4 mM CleanCap® AG reagent (TriLink® BioTechnologies, TRN7113), transcription buffer (40 mM Tris·HCl pH 8.0, 16.5 mM magnesium acetate, 10 mM dithiothreitol (DTT), 20 mM spermidine, 0.002 % (v/v) Triton X-100), 2 U/mL yeast inorganic pyrophosphatase (NEB®) and 1000 U/mL murine RNase inhibitor (NEB®). The IVT reaction was incubated at 37 °C for three hours and terminated by incubation with 200 U/mL NEB® DNase I at 37 °C for 15 minutes. The mRNA product was purified using a Monarch® RNA Cleanup Kit (NEB®) according to the manufacturer’s instructions, eluted in 1 mM sodium citrate and sterile filtered with a 0.22 μm syringe filter. The mRNA was quantified by UV spectrophotometry and integrity confirmed using the Agilent® TapeStation™.

### LNP production

For LNP formulation, a total lipid concentration of 15 mg/mL was used in the molar ratio of 50 SM-102 : 10 DSPC : 38.5 cholesterol : 1.5 DMG-PEG2000. The lipid mixture was made up to 3.75 mL with molecular grade 100 % ethanol. 1.5 mg of purified mRNA were combined with 11.25 mL of 0.1 M sodium acetate, pH 4.0. Formulation was performed on the NanoAssemblr® Ignite™ platform with the following parameters: total volume 13.5 mL, total flow rate 12 mL/min and flow rate ratio (aqueous:organic) 3:1. The mRNA-LNPs were dialysed using the Slide-A-Lyzer™ dialysis cassette (10K MWCO) and concentrated using an Amicon® Ultra-15 Centrifugal Filter Unit (10 kDa MWCO). The concentrated mRNA-LNPs were filtered with a 0.22 μm syringe filter and 0.2 volumes of 50 % sucrose were added by gentle pipetting for a final 10 % concentration.

### Bispecific antibody production

Bispecific antibodies were produced as previously described^34^. An scFv specific for PEG was linked to an scFv specific for human epidermal growth factor receptor^35^ or folate hydrolase 1^36^ via a glycine serine linker (G4S). The BsAb sequences were codon optimized for expression in C. griseus cells, included a κ light chain leader sequence for protein secretion, a 6 × Histidine motif at the N-terminus of the BsAb and a c-myc epitope tag at the C-terminus for purification and detection of the BsAb. The BsAb genes were cloned into the pcDNA™ 3.1 (+) mammalian expression plasmid (Invitrogen™) using HindIII and NotI restriction sites. For transient transfection, the plasmid DNA was transfected into ExpiCHO™ cells (Gibco™) using 2 μg DNA per mL cells at a concentration of 6 × 10^6^ mL^−1^ cells. For a 200 mL cell volume transfection, 200 μg DNA in 8 mL of OptiPRO™ serum free medium (SFM; Gibco™) were mixed with 7.4 mL OptiPRO™ SFM containing 640 μL ExpiFectamine™ (Gibco™) in OptiPRO™ SFM for five minutes prior to transfecting ExpiCHO™ cells. The transfected cells were cultured in ExpiCHO™ expression medium (Gibco™) at 37 °C, 7.5% CO2, 70 % humidity, with shaking at 130 rpm for 24 h, before feeding with 10 % ExpiCHO™ Feed (Gibco™) and 1.2 mL ExpiFectamine™ enhancer reagent (Gibco™) and returning cultures to 32 °C, 7.5 % CO2, 70 % humidity, with shaking at 130 rpm.

Following transfection, the cells were pelleted by centrifugation at 5250 × g for 30 minutes and the supernatant was collected and filtered through a 0.22 μm PES membrane (Sartorius®). The BsAbs were purified from the supernatant utilising a 5 mL HisTrap™ excel column (Cytiva™), eluting the protein with 20 × 10^−3^ M sodium phosphate, 500 × 10^−3^ M sodium chloride, and 500 × 10^−3^ M Imidazole pH 7.4. BsAbs were then buffer exchanged into 1 × phosphate-buffered saline (PBS) using a HiPrep™ 26/10 column (Cytiva™). The final product was sterile filtered using a 0.2 μm PES membrane filter (Sartorius®).

### Characterisation of LNPs

LNP size and charge were measured on a Zetasizer® Ultra (Malvern Panalytical®). mRNA-LNPs were diluted 50-fold in distilled water (Invitrogen™). For pre-mixing, 10 × excess BsAbs (w/w) were added and samples were incubated for 60 minutes at room temperature.

### Cell culture

MDA-MB-468 human breast cancer cells were cultured in Dulbecco’s modified Eagle’s medium (DMEM; Gibco™), supplemented with 10 % (v/v) fetal bovine serum (FBS; Gibco™) and 1 × penicillin-streptomycin (P/S; 100 U/mL penicillin and 100 μg/mL streptomycin; Gibco™). LNCaP human prostate cancer cells were maintained in Roswell Park Memorial Institute medium (RPMI-1640; Sigma-Aldrich®), 10 % FBS and 1 × P/S. Cells were incubated at 37 °C in 5 % CO2 and propagated for no more than 30 passages.

### *In vitro* BsAb-targeted mRNA-LNP delivery

For 70 % confluency at transfection, 1.9 × 10^5^ MDA-MB-468 or LNCaP cells were plated into the wells of 24-well cell culture plates in 500 μL of appropriate cell culture medium. To enhance LNCaP adhesion, 24-well plates were coated with poly-D-lysine (Gibco™) according to manufacturer specifications. The cells were allowed to attach overnight before treatment. All treatments were incubated with cells for 60 minutes at 37 °C with 5 % CO2. Treatments were diluted in Dulbecco’s Phosphate-Buffered Saline (DPBS; Gibco™) and added in volumes of 10 μL. mRNA concentration per well was 60 ng and BsAb concentration was 600 ng. For washing steps, cells were rinsed twice with DPBS and fresh complete medium was added. Four hours after addition of LNPs, cells were prepared for flow cytometry.

For pre-mixed samples, eGFP mRNA-LNPs were incubated with BsAbs for 60 minutes at room temperature. The treatment was then added to the wells, incubated and washed. For pre-targeting, cells were incubated with BsAbs for 60 minutes, washed, incubated with eGFP mRNA-LNPs and washed again. The untargeted LNP samples were incubated with DPBS, washed, incubated with eGFP mRNA-LNPs and washed again.

### Flow cytometry

To prepare samples for flow cytometry, medium was removed, cells were washed with DPBS, detached with 0.25 % 1 × trypsin (Gibco™), centrifuged at 200 × g for 5 minutes and resuspended in 250 μL flow buffer (DPBS, 2 % FBS, 2 mM ethylenediaminetetraacetic acid; EDTA). 7-aminoactinomycin D (7-AAD; Invitrogen™) viability stain was used and samples were incubated in the dark on ice for 30 minutes. 20,000 single cell events were recorded on a CytoFLEX™ Flow Cytometer (Beckman Coulter®) at a flow rate of 10 μL/s. eGFP and 7-AAD were excited using a 488 nm laser and emission was detected with a 525/40 bandpass and a 690/50 bandpass filter, respectively. Compensation and data analysis were performed using FlowJo™ v10.10.0. Software^37^ (BD Life Sciences™). GraphPad Prism™ v10.1.2^38^ (GraphPad Software) was used for statistical analysis and graphing. Statistical analysis was performed using two-tailed t-tests assuming equal variance with *p < 0.05, **p < 0.01, ***p < 0.001 and ∗∗∗∗p < 0.0001.

### Microscopy

4 × 10^4^ MDA-MB-468 cells or 6 × 10^4^ LNCaP cells were seeded on coverslips in 24-well plates and incubated overnight at 37 °C with 5 % CO2. For LNCaP cells, coverslips were coated with poly-D-lysine (Gibco™) according to manufacturer instructions. The cells were treated as described for *in vitro* BsAb-targeted mRNA-LNP delivery. To prepare the samples for confocal microscopy, the cells were fixed with 4 % paraformaldehyde (Novachem™) for 15 minutes at room temperature, washed three times with 1 × PBS (Gibco™), permeabilised and blocked with PBS buffer containing 0.1 % Triton X-100 (Sigma-Aldrich®) and 1 × bovine serum albumin (Sigma-Aldrich®) for one hour at room temperature. DAPI (BioLegend®) and Protein L-phycoerythrin conjugate staining (1:100; Cell Signaling Technology®) was performed for one hour at room temperature. The cells were washed three times with 1 × PBS and one time with Milli-Q® water before mounting on glass slides. Imaging was performed on a Zeiss LSM 710 inverted laser scanning confocal microscope at the Queensland node of the NCRIS-enabled Australian National Fabrication Facility (ANFF). Images were taken using a 63 × oil immersion objective for sequential scanning with excitation of DAPI, eGFP and phycoerythrin at 405 nm, 488 nm and 561 nm, respectively. FIJI^39^ was used for image processing.

Cryo-transmission electron microscopy was performed by the Centre for Microscopy and Microanalysis, UQ.

### *In vivo* imaging of mRNA-LNP biodistribution

All studies were in accordance with guidelines of the Animal Ethics Committee of The University of Queensland (UQ; Approval 2023/AE000135), and the Australian Code for the Care and Use of Animals for Scientific Purposes. Female Balb/c nude mice (approximately 8 weeks of age) were acquired from the Ozgene-ARC (Western Australia) and housed in temperature and humidity-controlled housing with *ad libitum* access to food and water.

Mice were subcutaneously injected with 5 × 10^6^ MDA-MB-468 cells in the right flank (50 μL PBS, 27G needle). After tumours reached a palpable size (ca. 100-200 mm^3^), the mice were segregated into three cohorts of n = 3 animals where the targeting strategy was varied. Mice in the untargeted LNP group were injected intravenously via the lateral tail vein, with 10 μg of luciferase mRNA-LNPs diluted in 100 μL of PBS (29G needle). Mice in the pre-mixed cohort were injected intravenously with mRNA-LNPs pre-incubated with anti-PEG:anti-EGFR BsAbs at a ratio of 50 μg BsAbs per 10 μg mRNA 15 minutes prior to intravenous injection (diluted in 100 μL PBS, 29G needle). Mice in the pre-targeted cohort were injected intravenously with 1 mg of anti-PEG:anti-EGFR BsAbs (diluted in 100 μL of PBS with 300 mM NaCl) eight hours prior to injection of 10 μg of mRNA-LNPs diluted in 100 μL of PBS (29G needle).

Mice were injected intraperitoneally with 150 mg kg^-1^ of D-Luciferin (VivoGlo™) 15 minutes prior to imaging using an IVIS® Lumina™ X5 imaging system (PerkinElmer®) eight hours and 48 hours post injection of the LNPs. Data were acquired using the default acquisition settings, then post-processed to a binning factor of four. Segmented regions of interest (ROIs) were drawn to delineate the liver and tumour of mice for *in vivo* analysis and over the major clearance organs for *ex vivo* analysis. Background subtraction was performed using a mouse injected intravenously with PBS (100 μL PBS, 29G needle) and D-Luciferin delivered intraperitoneally as described above. Data were analysed using the Living Image® software (PerkinElmer®).

## Acknowledgements

We acknowledge the following sources of funding and support: Australian Government Research Training Program (RTP) Scholarship to B.D., Innovation Connections Funding to S.W.C. and T.R.M. National Health and Medical Research Council (GNT2019056) to K.J.T. and (GNT2014002 and GNT1161832) to T.R.M., Australian Research Council (IH220100017) to K.J.T and (DE230100036) to S.W.C., Medical Research Future Fund (MRFCRI000063) to S.W.C. and T.R.M. National Collaborative Research Infrastructure Strategy (NCRIS) to T.R.M. and S.W.C., Therapeutic Innovation Australia (TIA) to T.R.M., Tour de Cure to S.W.C., and The University of Queensland to S.W.C. and T.R.M. The authors acknowledge the facilities and the scientific and technical assistance of the Australian National Fabrication Facility (ANFF, Queensland Node), the Centre for Microscopy and Microanalysis (CMM), the National Imaging Facility (NIF) and BASE at The University of Queensland. BASE is supported by Therapeutic Innovation Australia (TIA). TIA is supported by the Australian Government through the National Collaborative Research Infrastructure Strategy (NCRIS) program.

## Author contributions

B.D. and J.H. performed the experiments and the analysis. C.B.H. and S.W.C. conceived the project. S.W.C., C.B.H., T.R.M. and K.J.T. funded the study. All authors contributed to writing the paper.

## Competing interests

T.R.M. and S.W.C. have received research funding from Oxford Nanopore Technologies®, Sartorius® Stedim Australia, and Sanofi ™. T.R.M. and S.W.C. have received support for conference attendance, travel and accommodation from Moderna® and Oxford Nanopore Technologies®. The other authors declare no competing interests.

**Supplementary fig 1:**
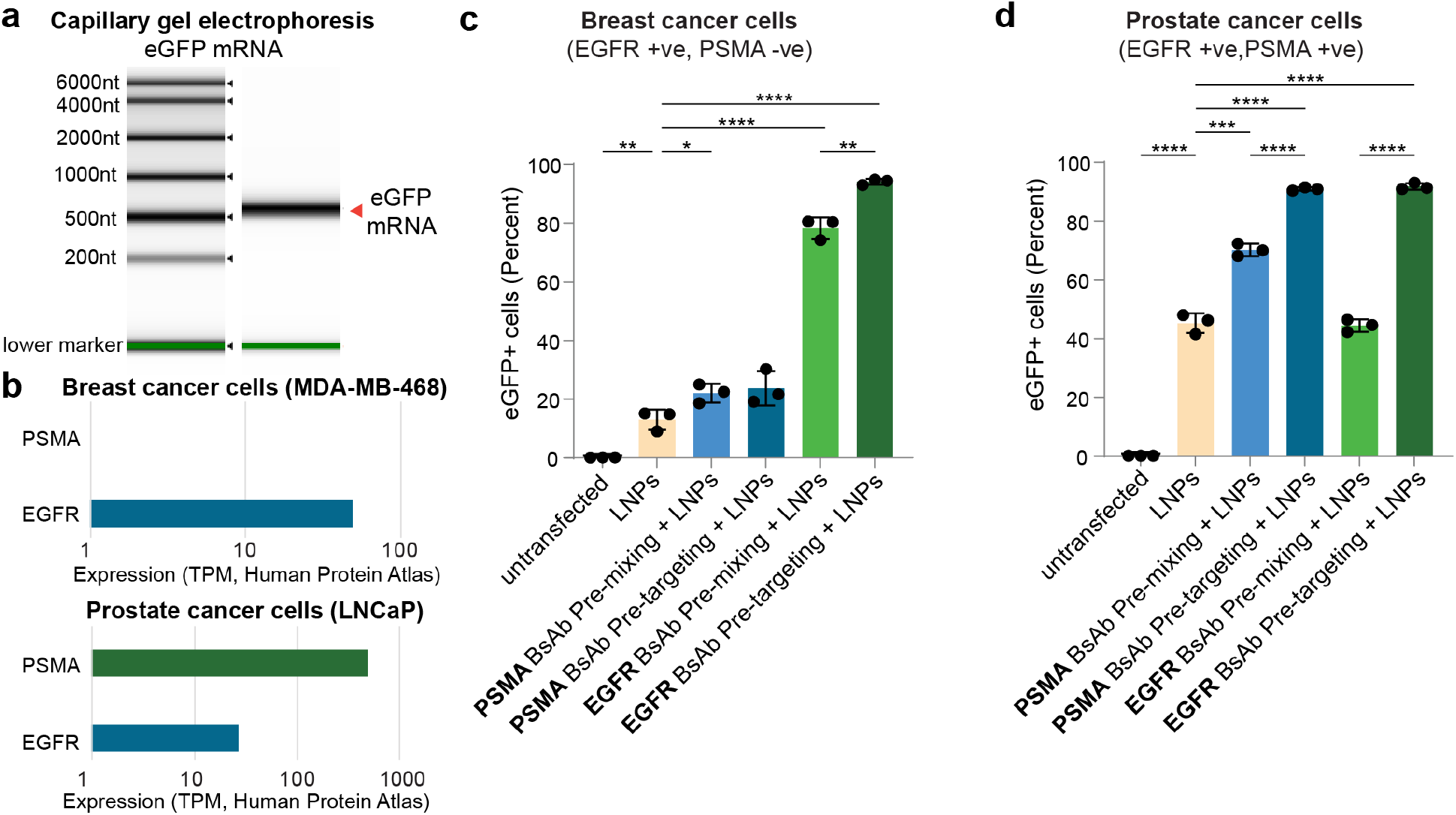
**a**, Analysis of size and purity of in vitro transcribed eGFP mRNA on electropherogram. **b**, Quantification of PSMA and EGFR RNA in MDA-MB-468 breast cancer cells and LNCaP prostate cancer cells, respectively, in transcripts per million (TPM; Human Protein Atlas proteinatlas.org). **c**, Percentage of eGFP expressing MDA-MB-468 breast cancer cells (EGFR+ve, PSMA-ve) transfected with eGFP-mRNA LNPs. EGFR-PEG BsAbs or PSMA-PEG BsAbs were pre-mixed with LNPs or pre-targeted to MDA-MB-468 cells, respectively. **d**, Percentage of eGFP expressing LNCaP prostate cancer cells (EGFR+ve, PSMA+ve) transfected with eGFP-mRNA LNPs. EGFR-PEG BsAbs or PSMA-PEG BsAbs were pre-mixed with LNPs or pre-targeted to MDA-MB-468 cells, respectively. eGFP expression was measured using flow cytometry. Statistical analysis was performed using two-tailed t-tests assuming equal variance. Bars represent the mean value, error bars indicate standard deviation (n = 3). *p < 0.05, **p < 0.01, ***p < 0.001 and ∗∗∗∗p < 0.0001.

**Supplementary fig 2:**
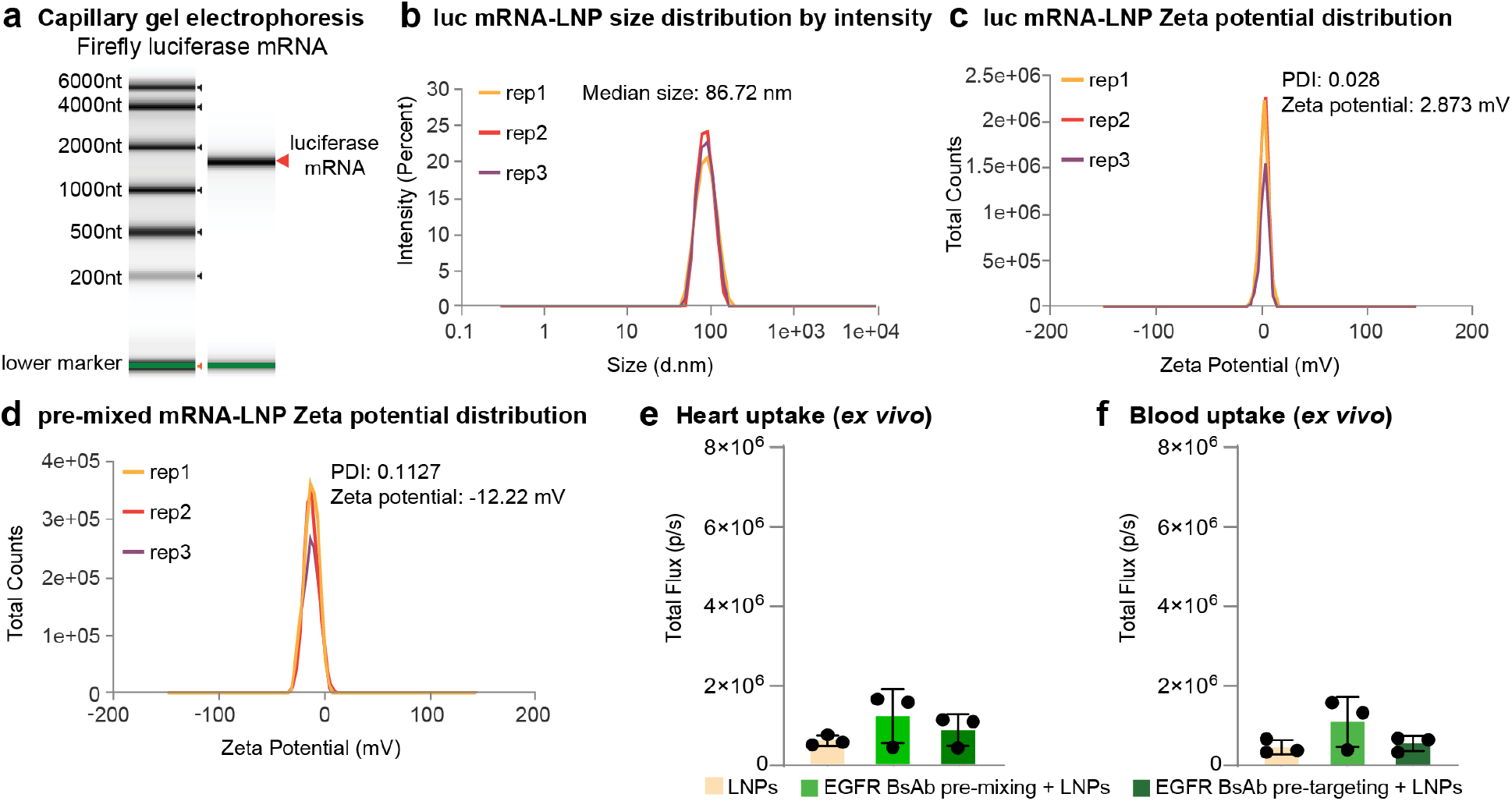
**a**, Analysis of size and purity of in vitro transcribed firefly luciferase mRNA on electropherogram. **b**, Triplicate dynamic light scattering measurements of size distribution of luciferase-mRNA LNPs. **c**, Triplicate electrophoretic light scattering measurements of zeta potential distribution of luciferase-mRNA LNPs. **d**, Triplicate electrophoretic light scattering measurements of zeta potential distribution of eGFP-mRNA LNPs pre-mixed with PSMA BsAbs. **e**, Ex vivo bioluminescence imaging of heart and **f**, blood. Background was subtracted based on a saline-injected mouse. Bars represent the mean value, error bars indicate standard deviation (n = 3).

